# Modeling conditional distributions of neural and behavioral data with masked variational autoencoders

**DOI:** 10.1101/2024.04.19.590082

**Authors:** Auguste Schulz, Julius Vetter, Richard Gao, Daniel Morales, Victor Lobato-Rios, Pavan Ramdya, Pedro J. Gonçalves, Jakob H. Macke

## Abstract

Extracting the relationship between high-dimensional recordings of neural activity and complex behavior is a ubiquitous problem in systems neuroscience. Toward this goal, encoding and decoding models attempt to infer the conditional distribution of neural activity given behavior and vice versa, while dimensionality reduction techniques aim to extract interpretable low-dimensional representations. Variational autoencoders (VAEs) are flexible deep-learning models commonly used to infer low-dimensional embeddings of neural or behavioral data. However, it is challenging for VAEs to accurately model arbitrary conditional distributions, such as those encountered in neural encoding and decoding, and even more so simultaneously. Here, we present a VAE-based approach for accurately calculating such conditional distributions. We validate our approach on a task with known ground truth and demonstrate the applicability to high-dimensional behavioral time series by retrieving the conditional distributions over masked body parts of walking flies. Finally, we probabilistically decode motor trajectories from neural population activity in a monkey reach task and query the same VAE for the encoding distribution of neural activity given behavior. Our approach provides a unifying perspective on joint dimensionality reduction and learning conditional distributions of neural and behavioral data, which will allow for scaling common analyses in neuroscience to today’s high-dimensional multi-modal datasets.

## Introduction

Recent developments in experimental techniques allow real-time behavioral tracking of animals (***Mathis et al., 2018***; ***Günel et al., 2019***; ***Pereira et al., 2019***), and simultaneous recordings of hundreds of neurons across multiple brain regions (***Ahrens et al., 2013***; ***Sofroniew et al., 2016***; ***Jun et al., 2017***). Modern datasets in neuroscience are thus increasingly large, high-dimensional (***de Vries et al., 2020***; ***Siegle et al., 2021***), and commonly consist of multiple modalities—e.g., neural activity and behavior (***Mimica et al., 2023***)—that often have highly non-linear relationships (***Sani et al., 2021b***). While data collection has changed drastically in the last years, an important goal of systems neuroscience remains the same: understanding how brain activity gives rise to complex behavior.

To gain insights from neural and behavioral data, neuroscientists have developed various neural encoding and decoding models (***Kriegeskorte and Douglas, 2019***). These tasks should be ideally addressed in a probabilistic manner to account for the inherent variability of neural and behavioral measurements and in order to quantify resulting uncertainty. As experimental neuroscience is moving towards less controlled, unconstrained multi-modal data collection, this aspect becomes even more relevant. Both probabilistic encoding and decoding tasks can, algorithmically, be boiled down to the task of calculating conditional distributions: Encoding studies in neuroscience involve calculating the conditional distribution of neural activity given behavior or other observations such as stimuli (***Pillow et al., 2008***). Conversely, for decoding analyses, one needs to calculate the conditional distribution over behavior, given neural activity (***Figure 1a***, left). Generating interpretable, accessible neuroscientific predictions from complex, high-dimensional data directly is very challenging (***Paninski and Cunningham, 2018***; ***Chen and Pesaran, 2021***), highlighting the need for tools that can infer low-dimensional representations of high-dimensional neural and behavioral datasets (***Figure 1a***, right). In short, to gain neuroscientific insights from such complex datasets, our goal is to unify 1) the ability to link neural and behavioral data (i.e., through encoding/decoding models, ***Figure 1a***, left), and 2) joint dimensionality reduction of the data, ideally in a probabilistic and generative manner (***Figure 1a***, right).

**Figure 1.**
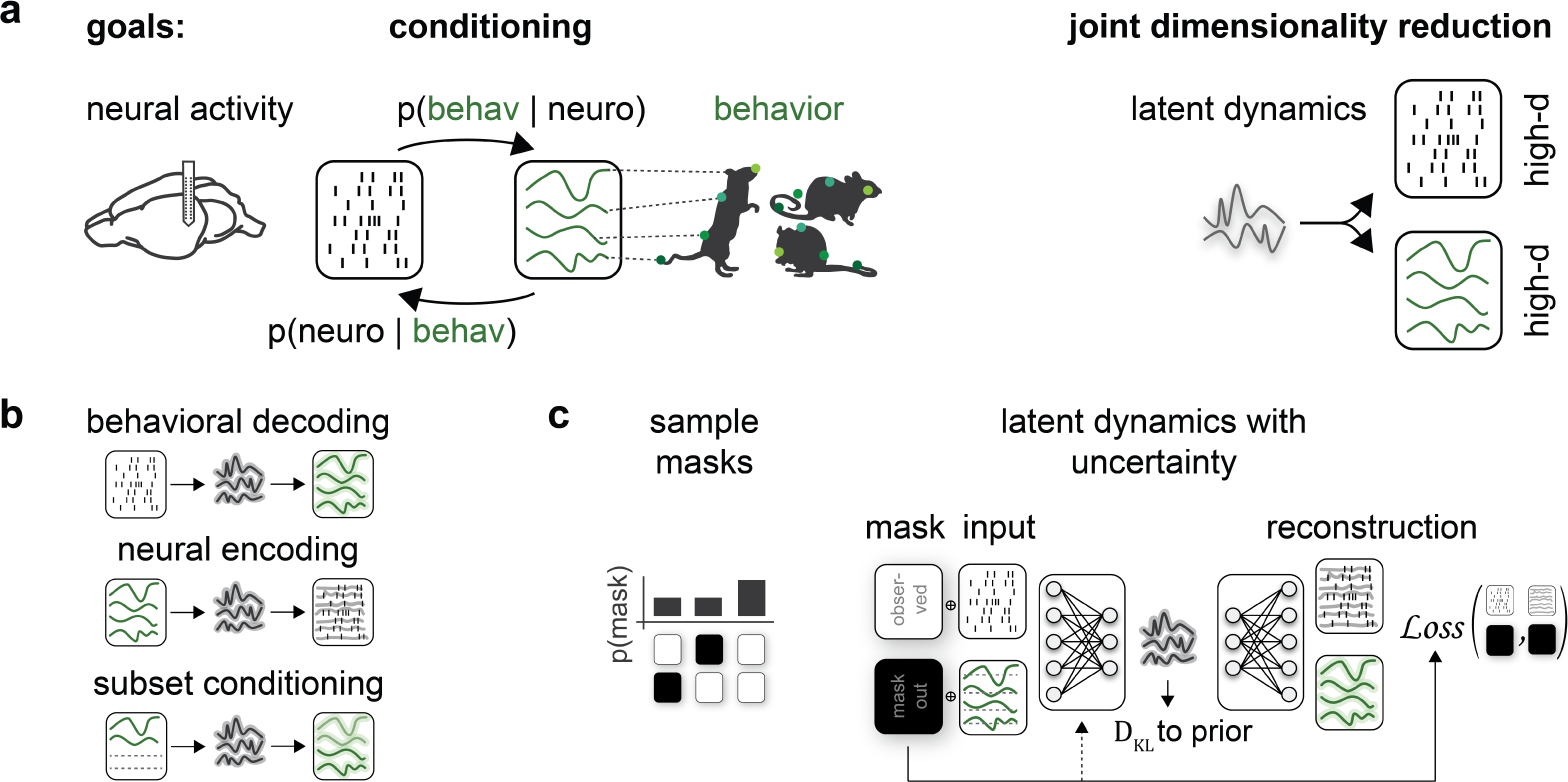
Latent variable models that can deal with conditional distributions arising in large-scale multi-modal datasets a Our approach aims to address two common goals of (neuro)scientific analyses: Conditioning, e.g., for linking brain activity (neuro) and behavior (behav) and joint dimensionality reduction of potentially high-dimensional neural and behavioral data. b Conditional distributions arise, e.g., when learning the distribution over behavior given brain activity (behavioral decoding), the distribution over neural activity given behavior (neural encoding), or when analyzing the interaction of data subsets such as different behavioral variables or multiple brain regions. c Masked variational autoencoder training scheme for data of potentially different data types (e.g., count and continuous) with structured masks for modeling conditional distributions. During each training iteration, one mask is chosen randomly. The reconstruction loss is computed solely on observed data. The Kullback-Leibler divergence *D*_*KL*_ of the inferred latent representation and a defined prior leads to a regularization of the latent space (see Meth- ods).

Various dimensionality reduction methods have demonstrated that a substantial fraction of variability both in unconstrained behavior and neural population activity can be captured by a few latent (i.e., unobserved) dimensions (***Yu et al., 2009***; ***Sussillo et al., 2016***; ***Batty et al., 2019***; ***Keeley et al., 2020***; ***Sani et al., 2021a***; ***Luxem et al., 2022***; ***Schneider et al., 2023***). This insight has driven the development of various latent variable models that infer underlying low-dimensional representations from neural data (***Pfau et al., 2013***; ***Sussillo et al., 2016***; ***Schimel et al., 2021***; ***Jensen et al., 2021***; ***Bashiri et al., 2021***). Classical methods often require simplifying modeling assumptions, such as (switching) linear dynamics (***Macke et al., 2011***; ***Petreska et al., 2011***; ***Linderman et al., 2017***), or strictly Gaussian observations (***Yu et al., 2009***), and rely on model-specific optimization schemes, e.g., Expectation Maximization algorithms or subspace-identification methods (***Buesing et al., 2012***; ***Sani et al., 2021a***). With the rise of deep learning and more flexible optimization schemes, various of these assumptions can be relaxed (***Sussillo et al., 2016***; ***Luxem et al., 2022***; ***Schneider et al., 2023***), leading to latent variable models that can capture complicated non-linear relationships in the data and underlying low-dimensional dynamics (***Girin et al., 2021***).

One such model class, exploiting deep inference networks, is the variational autoencoder (VAE) (***Rezende et al., 2014***; ***Kingma and Welling, 2014***). Inference networks of VAEs take observed data as the input and return a distribution over the latent state. Sequential variants of VAEs can infer latent representations underlying heterogeneous time-series datasets, e.g., consisting of both continuous and count data (spiking) (***Nazábal et al., 2020***; ***Girin et al., 2021***; ***Brenner et al., 2024***) and are commonly used in analyzing neural and behavioral data (***Sussillo et al., 2016***; ***Zhou and Wei, 2020***; ***Schimel et al., 2021***; ***Luxem et al., 2022***). However, most VAE-based methods cannot adequately deal with a ubiquitous analysis task in neuroscience: calculating arbitrary conditional distributions *p* (data subset A ∣ data subset B) (***Williams et al., 2019***; ***Ivanov et al., 2019***; ***Collier et al., 2020***). The reason for this is that inference networks of VAEs typically can only deal with *fully* observed input data, and have no means to model the (additional) uncertainty arising from partial observations.

Such conditional distributions do not only arise when estimating behavioral decoding (***Figure 1b***, top) and neural encoding distributions (***Figure 1b***, middle) but also become relevant when dealing with partially observed data or when studying interactions between brain regions or tracked body parts (***Figure 1b***, bottom). An ideal model should correctly estimate how uncertain it is about the inferred underlying representation and its predictions (error bands, ***Figure 1b***). Accurate uncertainty estimates can tell us how constrained one subset is given the other subset and let us reason about their dependencies beyond accuracy or similarity scores. However, in neuroscience, the quality of uncertainty estimates is typically not assessed.

In this work, we present an approach that enables VAEs to accurately model arbitrary data-conditionals arising in neuroscience. Specifically, we use a masked-training approach and demonstrate it in a variety of neuroscience applications where we achieve both dimensionality reduction and sampling of conditional distributions. Furthermore, we propose calibration tests to assess the quality of the generated conditional distributions.

VAEs have been extended to model the distribution of a missing data subset given an observed data subset (***Ivanov et al., 2019***; ***Nazábal et al., 2020***; ***Collier et al., 2020***). Conceptually, we build on these results and treat the modality we want to learn the conditional distribution over as missing. For example, to capture a behavioral decoding distribution, we modify the loss and training scheme of a VAE during joint training on neural and behavioral data by stochastically masking behavior. Our training approach is not limited to a specific modeling architecture but can be applied to a variety of VAE approaches. We showcase our approach on diverse datasets: sequential and

static, multi- and uni-modality, discrete and continuous datasets, each of which with a different VAE architecture. We first validate our approach on a task on which we have access to the ground-truth distributions. We demonstrate that our approach allows for correct inference of low-dimensional latents and accurate predictions. On a high-dimensional behavioral dataset of walking flies, we successfully recover the relationship between different body parts along with uncertainties and obtain realistic samples from the conditional distributions of masked legs. Finally, we showcase the approach in a challenging multi-modal neural and behavioral dataset, where we model encoding and decoding distributions of high-dimensional population activity from primary motor areas and self-paced reach movements (***O’Doherty et al., 2017***).

## Results

### Masked training of variational autoencoders for estimating conditional distributions

To prepare variational autoencoders to deal with conditional distributions commonly arising in neuroscience, we modify the training scheme of classical VAEs. During joint training on multiple data subsets, our approach prepares the network for each subset to be *structurally masked* at test time (***Figure 1c***). We use the term *structured masking* to refer to algorithmic masking of data subsets for conditioning to avoid confusion with actual data missingness—e.g., individual input channels that drop in and out. First, we specify the *structured masks* depending on the desired conditional distributions and specify how often (on average) each mask should be selected during training (***Figure 1c***, left, see Methods). We calculate the reconstruction loss solely considering *observed* subsets (see Methods) in the evidence lower bound, which is the optimization target of VAEs (***Kingma and Welling, 2014***; ***Collier et al., 2020***). To make it easier for the network to learn that an input has been masked, one can additionally pass the binary mask to the encoder network.

In summary, we propose modeling conditional distributions with VAEs, e.g., for neural encoding and behavioral decoding, by recasting it as a *structured masking* problem. This approach allows us to sample from a distribution of interest, e.g., to visualize time series of various potential behaviors that are likely given a neural population activity trace and vice versa. The generality of this approach allows for applying it to a variety of conditional distributions and variational autoencoder settings.

### Inference of conditionals in a tractable Gaussian Latent Variable Model (GLVM)

First, we evaluated if our training scheme and loss modification allow us to learn the correct distributions of interest on a simulated dataset where we have access to the ground-truth conditional and posterior distributions. This dataset was generated from a Gaussian Latent Variable Model (GLVM) with latent (unobserved) random variable *z* and data dimensions *x*, which linearly depend on the latent *z* (***Figure 2a***, see Methods). In this illustrative example, we can think of a subset of *x* as the high-dimensional neural activity, another subset of *x* as high-dimensional behavior, and the latent variable *z* as the low-dimensional representation underlying both neural and behavioral data. The inference network infers the distribution over these unobserved latents given a chosen *x*—i.e., it calculates the posterior distribution *p* (*z* observed x)— effectively inverting the data generation process in a probabilistic way. The strength of the linear coupling and the noise levels of individual *x* dimensions define how much information about the latent can be gained by observing those *x* dimensions.

**Figure 2.**
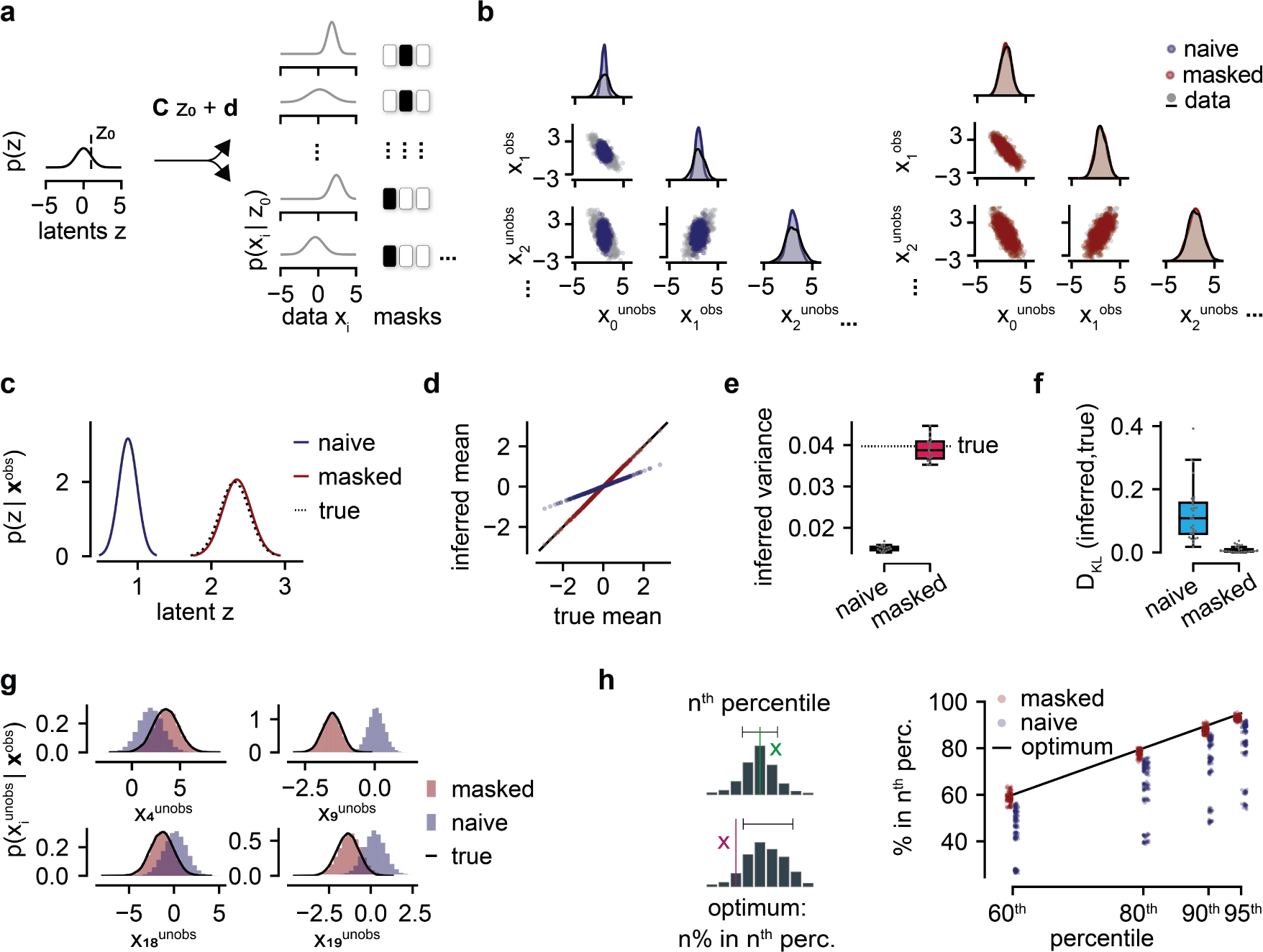
Inference of conditionals in a Gaussian Latent Variable Model (GLVM). a Left: Schematic of a GLVM with fixed parameters *θ* = *C*, *d*, Λ, where Λ = diag(σ_*i*_^2^) for *i* ∈ {1, 20}. Right: Structured masks for conditioning specified by the user. Here, with either 50% of the values masked or fully observed (0% masked). b-h Comparison of our masked training approach (red) with a vanilla VAE approach (blue, naive) when half of the inputs *x* are masked out at test time. b Model reconstructions and true test data samples (1D and 2D marginal distributions over *x*). c Inferred and analytical (true) posterior distributions over latent z given only the observed dimension of one test sample, i.e., *p*(*z* ∣ **x**^obs^). d True versus inferred posterior mean given a range of test samples. e Inferred posterior variance across multiple instantiations (seeds) and true posterior variance (dotted line). Box plots show median, minimum, maximum, 1st quartile, and 3rd quartile. f Average Kullback-Leibler divergence between true and inferred posterior distributions across different GLVM parameters (*θ* = *C*, *d*, Λ) and structured masks. g Conditional distributions over randomly chosen masked *x* dimensions (see Methods) for the same test sample as in panel c. h Left: Schematic of statistical calibration, evaluating the quality of uncertainty estimates. Optimal calibration: n% of true data points lie in the nth percentile confidence interval of the sampling distribution. Example of an *x* within the interval (green, top) and one outside of it (red, bottom). Right: Calibration checks of predicted conditional distributions *p*(*x*^unobs^ ∣ **x**^obs^) for all masked *x*-dimensions across multiple model seeds.

We contrast the masked VAE with a regular VAE trained on all data (referred to as naive training) regarding the capacity to capture data-conditionals *p* ( masked x observed x ) at test time (***Figure 2***). The naive approach fails to capture the true data distribution (grey), with overly narrow 1D and 2D (marginal) distributions (***Figure 2b*** left). Masked VAEs, however, can successfully reconstruct observed values (*x*_*i*_^*obs*^) and impute masked ones (*x_i_*^*unobs*^) (***Figure 2b***, right). The reason for this discrepancy lies in the ability to learn the distribution over the latents (posterior distribution) when some of the input data are unobserved. Masked training can perfectly infer the true (analytically calculated) posterior. In contrast, naive training fails to do so (***Figure 2c***). The naive network does not detect masked values as such. Hence, its posterior mean inference is biased (***Figure 2d***), and the posterior variance is too small (***Figure 2c***). In short, naive VAE training leads to confidently wrong predictions, while the masked network correctly adjusts the uncertainty about its predictions. This finding generalizes across different parameter sets (*C*, *d*, *λ*) and masking conditions (***Figure 2f***, ***Figure S1***).

One of the advantages of VAEs is that once trained, sampling from VAEs is straightforward. Thus, we can easily investigate whether the conditional samples for unseen test data correspond to the conditional distribution we set out to model. Wrongly inferred posterior distributions will likely result in inaccurate conditional samples.

Indeed, for four example masked data dimensions (*x*^*unobs*^), the distribution of conditional samples from the naive VAE is wrong, whereas the masked training distribution matches the true (analytical) conditional (***Figure 2g***).

In real neuroscientific datasets, we do not have access to the true conditional distributions for comparison. For such cases, we propose to evaluate the quality of the inference method and its uncertainty estimates with an adaptation of simulation-based calibration (***Cook et al., 2006***; ***Talts et al., 2018***). These calibration checks allow us to also evaluate the quality of the uncertainty estimates, and thus go beyond the evaluation of mean squared error or log-likelihood of *structurally masked* values, which informs only about the quality of the mean predictions. More concretely, calibration checks count how often masked test sample values *x*^*unobs*^ lie in the respective predicted conditional sampling distribution (***Figure 2h***, see Methods). We found that for the masked training, predictions are well-calibrated, i.e., neither overconfident nor underconfident (values close to the diagonal ***Figure 2h***, red). In contrast, the naive approach is confidently wrong for many test cases (***Figure 2h***, blue). Overall, these results suggest that masked training allows us to infer both the correct posterior *p* (*z* only observed x ) and conditional distributions *p* (masked x only observed x) in a tractable, well-specified example. The results demonstrate the statistical challenges that arise when one aims to use VAEs to perform conditioning and provide a theoretical basis for applying the masked training approach to neuroscientific data.

### Probabilistic conditional modeling of masked keypoint trajectories in fly walking behavior

Next, we wanted to investigate if the masked approach is applicable to complex time-series data in neuroscience and can successfully model conditional distributions of scientific interest. In particular, we focused on an experiment that characterizes the (backward) walking behavior of the fruit fly, *Drosophila melanogaster*, and applied our masked training approach to a sequential VAE developed for this high-dimensional behavioral dataset.

We obtained the dataset by tracking the centroids of individual flies and aligning the video frames such that fly heads are all pointing upward (***Figure 3a***, left). We tracked 32 body parts (*x*, *y* each) with DeepLabCut (***Mathis et al., 2018***), resulting in a 64-dimensional time series (see Methods). To account for the temporal structure of the data, the VAE’s architecture has both convolutional elements and recurrent neural networks based on Gated Recurrent Units (***Cho et al., 2014***), as well as elements for non-linear dimensionality reduction (see Methods). Analogous to the GLVM case, we adapted the masked training scheme for this sequential VAE to allow for modeling the conditional distribution over a subset of the fly body keypoints, given the remaining ones (***Figure 3a***, right, masked legs in green, see Methods). Here, we chose to mask body keypoints that are crucial for walking behaviors and show characteristic variability during walking: hind claw, hind tibia-tarsal joint, mid tibia tarsus, and mid claw of the left side.

**Figure 3.**
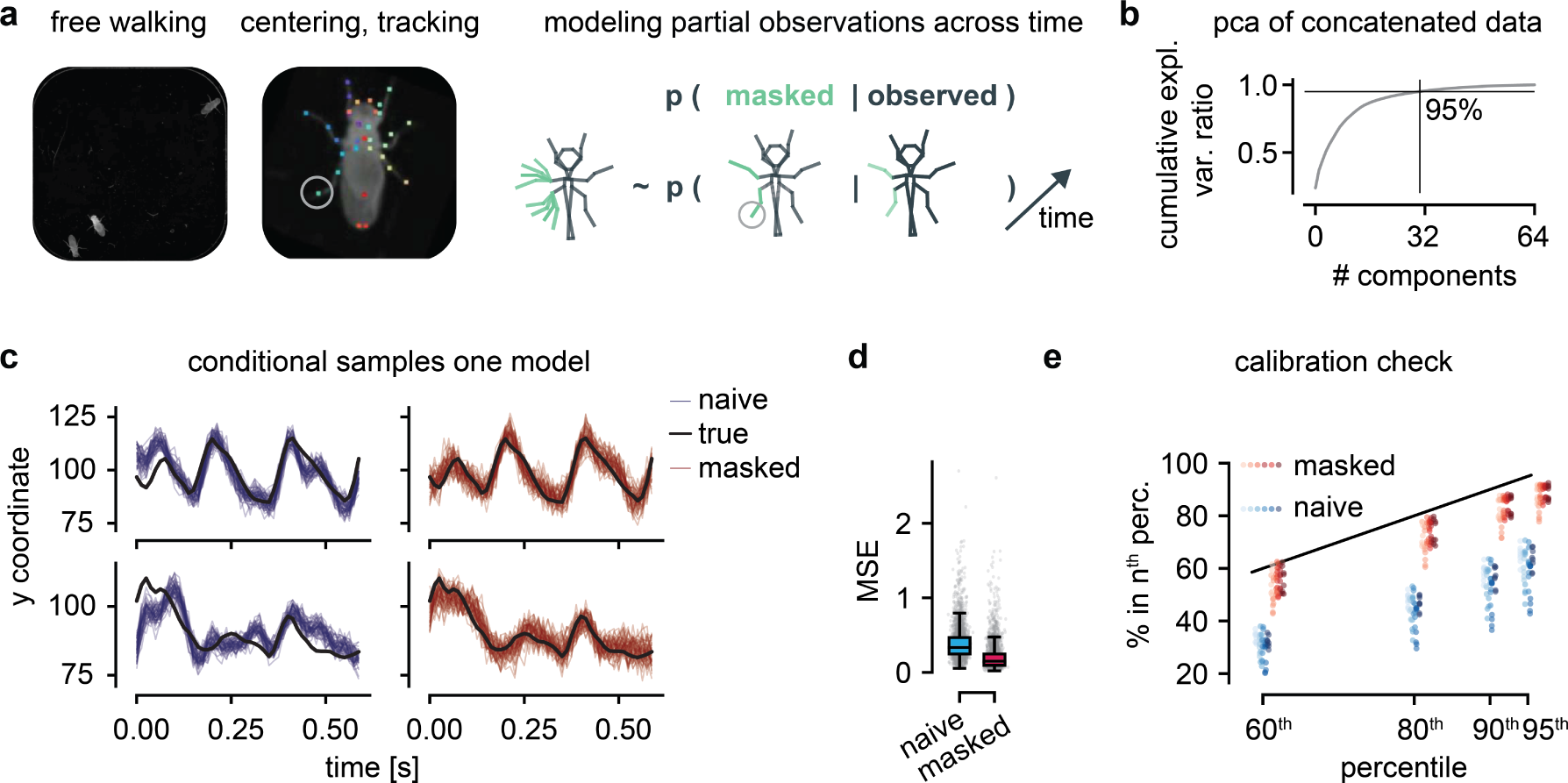
Conditional sampling of masked legs of walking flies. a Three flies are filmed from the bottom during walking behavior in a constrained arena. Cropped video frames of individual flies are centered, aligned for constant head direction, and tracked with Deeplabcut, resulting in a 64-dimensional time series. Schematic of the target distribution: the conditional distribution over masked left legs given the remaining body keypoints. b Cumulative explained variance of the number of components when performing principal component analysis on all concatenated time-series data. c Two example time series (from the test set) of the unobserved limb, marked with a circle in panel a. Conditional samples from the masked model in red, naive in blue, and true limb trajectory in grey. d Mean squared error (MSE) of mean predictions for the masked limb, averaged across time and test samples, for both training schemes. Box plots show the median and lower and upper quartiles. e Calibration checks of predicted conditional distributions *p* (limb^unobs^ limbs^obs^) for all masked leg keypoints shown in panel a, across multiple model seeds (different hue per seed). Optimal calibration in black.

If the model captures the dependence correctly via the compact latent representation, we expect accurate conditional modeling of these masked legs. In the previous example, the variability in the data was captured by one latent variable, but for experimental data, the underlying dimensionality is unknown. Here, we are dealing with a dataset with high intrinsic dimensionality: almost 32 principal components are required to capture 95% of the variance of the 64-dimensional dataset (***Figure 3b***). In our sequential VAE, we can exploit temporal dependencies and thus further reduce the dimensionality of the latent space while capturing stereotyped walking behavior (see Methods). Samples from the naive model capture overall trends of the masked left hind claw (circle in ***Figure 3a***) well, particularly during highly periodic walking (***Figure 3c***, left). However, ground-truth trajectories often deviate from the sampled trajectories (blue). In contrast, masked training produces more faithful predictions and uncertainty estimates (red), reflected in the inclusion of the ground truth for most time steps (***Figure 3c***, right). This is also captured by a lower average mean squared error (MSE) of the masked VAE test predictions (averaged across time) of the masked keypoint shown in panel c. Yet, MSE alone does not immediately reveal a substantial performance boost through masked training (***Figure 3d***). The difference, however, becomes clear when inspecting the uncertainty estimates: when the naive approach is wrong, it is confidently wrong (large deviations from the diagonal in ***Figure 3e***), while the masked approach is better calibrated.

We conclude that our masked training methodology is indeed applicable to time-series datasets and allows us to faithfully model the conditional distributions of masked body keypoints given the remaining ones. Masked training leads to better uncertainty estimates—it allows the network to know better when it does not know.

### Decoding continuous reaches from neural population activity

To be effective for neuroscientific research, our method should be able to deal with data types and modalities that commonly arise in neuroscience. Therefore, we implemented the masked training scheme for a classic monkey reach task, which is particularly challenging due to its continuous instead of trial-based structure (using publicly available data from ***O’Doherty et al.*** (***2017***), ***Figure 4a***, left). We focus on the behavioral decoding distributions, i.e., the conditional distribution of *x*- and *y*-reach directions given activity traces of >200 neurons (***Figure 4a***, right).

**Figure 4.**
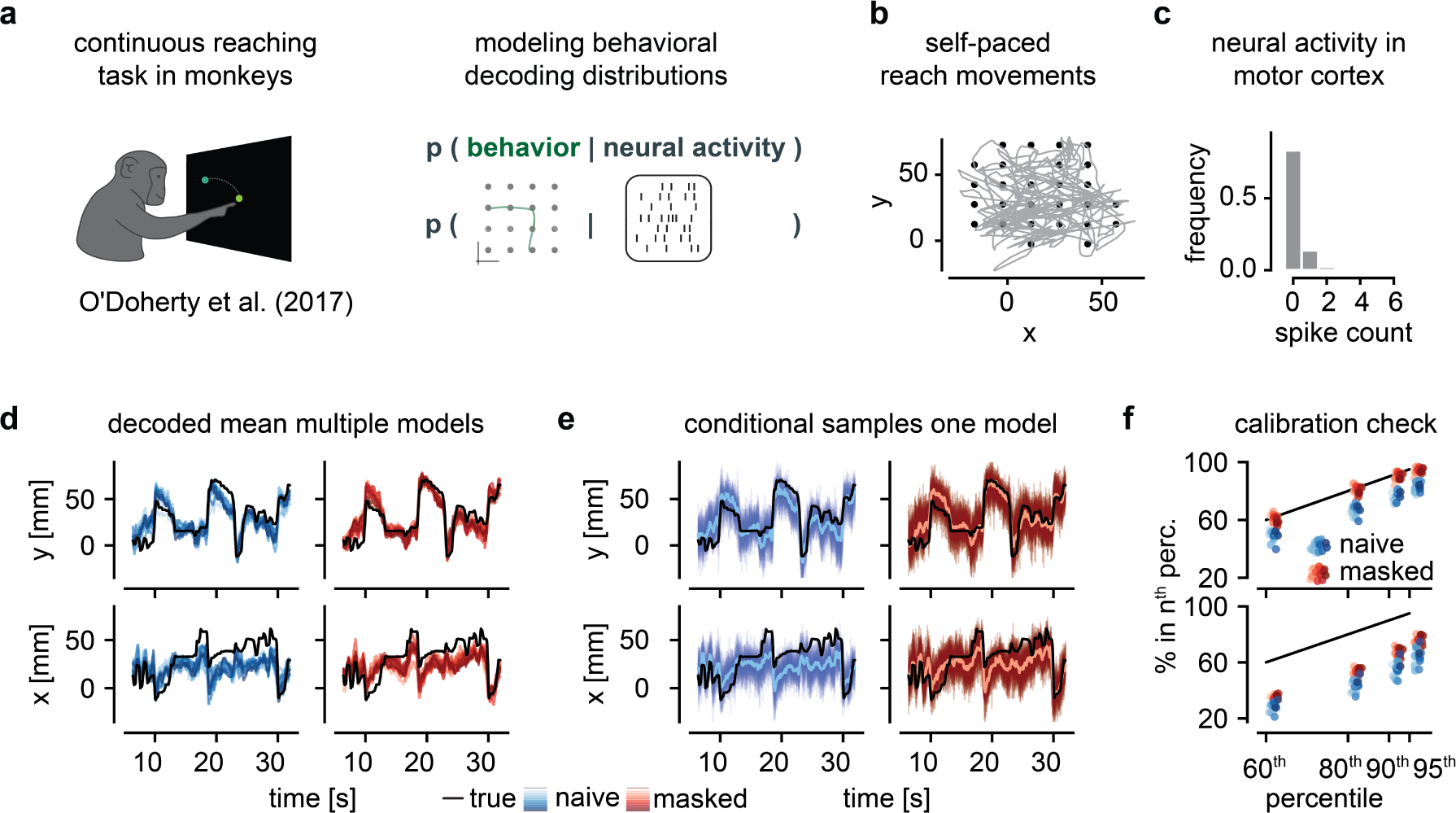
Behavioral decoding of continuous reach movements from monkey primary motor cortex a A monkey performs self-paced, continuous reaches on an 8x8 grid with simultaneous cortical electrophysiology recordings. Schematic of the target distribution: the conditional distribution over behavior, in this case, cursor trajectories given neural population activity recorded in monkey primary motor cortex. b Behavioral trace of continuous reach movements (grey) and targets (black). c Frequency of observed spike counts across primary motor cortex during the reach movements shown in panel b, binned at 64ms. d Example traces of cursor positions and the mean predictions for the naive (blue) and masked (red) modeling approaches for multiple model seeds (different hue per seed). e Same cursor traces as in panel d but with conditional samples from a single naive (blue) and masked (red) model, respectively. f Calibration checks of predicted conditional distributions *p* (behavior neural activity) respectively for x-y-directions, across multiple model seeds. Optimal calibration in grey. d-f The y-direction is in the top and the x-direction in the bottom row.

The monkey reaches toward an indicated light target on an 8x8 grid, leading to movements of different lengths, het- erogeneous angles, and velocities (***Figure 4b***). Neural activity is simultaneously recorded in primary motor cortex.

The maximum spike count of individual units is six spikes in time windows of 64 ms (***Figure 4c***, binning consistent with ***Makin et al.*** (***2018***) to capture behaviorally relevant timescales). We built a sequential VAE for this multi-modal dataset, which consists of both continuous (behavioral) and discrete data (spike counts). Our reconstruction loss, therefore, is composed of a Poisson- (for spike counts) and Gaussian- (for behavior) negative log-likelihood (GNLL) loss (see Methods, ***Nazábal et al.*** (***2020***); ***Brenner et al.*** (***2024***)). We specified masks for encoding and decoding distributions during training, i.e., we masked either neural activity (replaced by zeros) or behavior (replaced by the mean cursor position).

Masked mean reconstructions (red) of behavioral traces are more accurate than naive (blue) predictions across many model seeds (***Figure 4d***). Surprisingly, the naive approach is performing relatively well, and while we see some sections where the masked approach is performing better, the errors are quite consistent across the two approaches. This indicates that some sections of the traces are less correlated with neural activity than others, and both models are capable of exploiting some correlations required for conditional modeling. Sampling from individual masked and naive models (***Figure 4e***) and calibration checks (***Figure 4f***) again demonstrate that the masked but not naive approach targets the conditional—the decoding distribution. Note that a discrepancy between *x* (bottom rows) and *y* (top) decoding performance (***Figure 4d-f***) has been reported previously for this dataset (***O’Doherty et al., 2017***; ***Makin et al., 2018***; ***Pei et al., 2021***). While the masked VAE is not perfectly calibrated on this dataset, it clearly outperforms the naive VAE.

In conclusion, our masked training approach can be readily applied to multi-modality datasets and makes it possible to sample from a conditional distribution over a continuous time series given high-dimensional time series of discrete count data.

### Encoding of continuous reaches in neural population activity

Next, we assessed the performance of the same trained model on the reversed and more challenging task: modeling the high-dimensional conditional distribution over the activities of 213 neurons in primary motor cortex given only the two-dimensional behavioral trajectories (***Figure 5a***).

**Figure 5.**
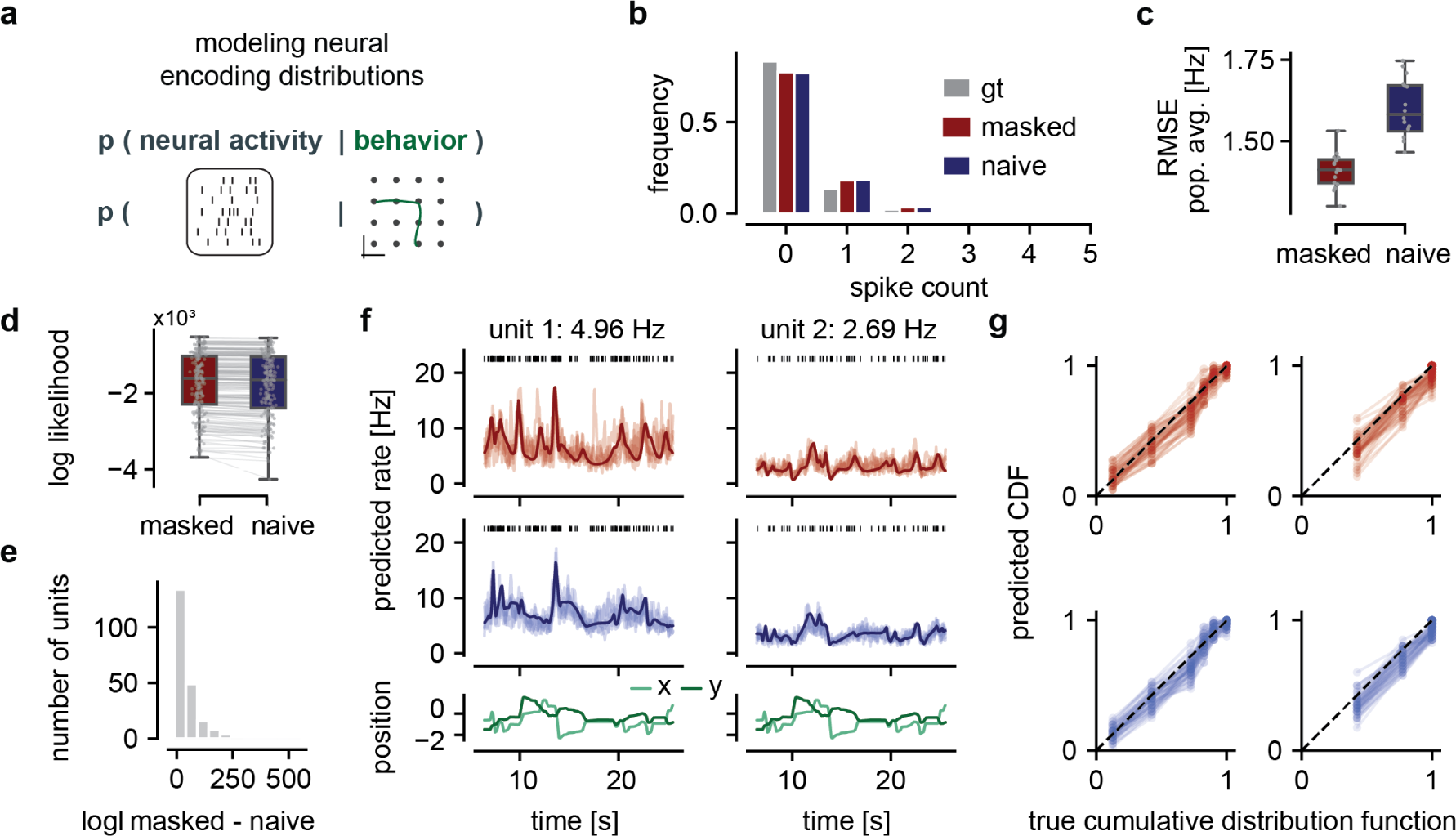
Neural encoding distributions given the continuous movements in the monkey reach task a Schematic of the target distribution: the conditional distribution over neural population activity recorded in monkey primary motor cortex given behavior, i.e., cursor trajectories. b Frequency of observed spike counts across primary motor cortex during reach movements binned at 64ms and the predicted spike count distribution of the masked (red) and naive (blue) approach. c Root mean squared error (RMSE) of the population average and predicted population average averaged across time for different model seeds. d Log-likelihood per neuron for the masked and naive approaches averaged across time points and model seeds. Higher is better. e Distribution of differences in log-likelihood (logl) of the masked minus the naive approach. Positive values indicate a better model fit of the masked approach. f Sampled rate predictions (10 each) and mean rate prediction from both models (masked red top, naive blue middle) for two example neurons with different activity levels given the standardized behavioral trajectory (bottom). g Cumulative distribution function of the observed spikes (true) vs. predicted spike distributions sampled from the masked (red) and naive (blue) VAE for the rate predictions shown in panel f. To calculate the CDF, spike counts are aggregated across five bins due to low spike counts. Optimal predictions would lie on the diagonal (black dotted line).

Both masked and naive approaches generally capture the histogram of observed spike counts given the reach trajectories (***Figure 5b***) – despite a slight over-prediction of spike counts –, but the naive approach has worse population average estimates across model instantiations (***Figure 5c***). The log-likelihood per neuron across model instantiations reveals the superior performance of the masked versus naive encoding (***Figure 5d,e***).

Samples from the trained models suggest that both masked and naive approaches correctly predict time-varying firing rates that clearly reflect the reach movements (***Figure 5f***). Notably, the masked approach reveals higher variability in the rate predictions reflecting higher posterior uncertainty.

Spike counts are discrete rather than continuous variables, so we adapted the method to assess the uncertainty calibration: we compare the cumulative distribution function (CDF) of the ground-truth spike train against a Poisson spike train with a rate sampled from either the masked or the naive models (***Figure 5g***). We find that the sampling distribution for both masked and naive capture the ground truth distribution reasonably well (***Figure 5g***; for other units see appendix).

In conclusion, our masked VAE approach allows us to model and produce samples from both decoding and encoding distributions in one single model without requiring any retraining.

## Discussion

We introduce a training methodology for modeling conditional distributions with masked variational autoencoders, bridging dimensionality reduction, generative modeling, and encoding and decoding analyses in neuroscience. Our experiments show that modifying the training scheme and loss through *structured masking* enables VAEs to model specified conditional distributions. Thus, our approach allows for joint dimensionality reduction of high-dimensional multi-modal data and conditioning on specified modalities. It is not restricted to specific architectural or modeling choices and can be easily applied to a variety of variational autoencoders deployed in neuroscience. We validated our approach on a tractable example in which we correctly learned the ground-truth posterior and conditional distributions. We applied our approach to two neuroscientific time-series datasets: a continuous reach task in monkeys (***O’Doherty et al., 2017***), in which we probabilistically encoded behavior in—and decoded behavior from—high-dimensional neural activity, as well as a behavioral dataset of walking flies, for which we successfully modeled the conditional distributions of masked body parts. Furthermore, we showed how to assess the models’ uncertainty estimates, a crucial but often neglected aspect in deep-learning-based dimensionality reduction, and verified that our models learn calibrated distributions.

### Generality of modeling conditional distributions in neuroscience and beyond

A key contribution of this work lies in linking conditional distributions of neural and behavioral encoding and decoding to probabilistic approaches for dealing with missing data (***Nazábal et al., 2020***; ***Collier et al., 2020***).

The generality of this approach opens up various possibilities beyond encoding and decoding, as generating conditional samples and performing inference from partial observations has many applications. In experiments such as the fly walking example where occlusions or tracking issues occur, our approach enables sampling from distributions over the obscured body keypoints. In addition, the inherent denoising property of VAEs can correct noisy markers, especially when the observation noise is learned explicitly. More generally, data imputation can be framed as modeling conditional distributions over missing variables given observed ones. This is a relevant pre-processing step for many downstream analyses that require complete datasets in neuroscientific and clinical applications, as well as in other domains (***Talukder et al., 2022***; ***Vetter et al., 2023***). For example, if an electrode breaks during a neural recording, a masked VAE approach can salvage the dataset by computing conditionals for the failed electrode using complete data from other sessions.

### Using and assessing uncertainties in deep-learning based models

It is well-established that both modern deep learning models (***Guo et al., 2017***) and traditional Bayesian decoders (***Wei et al., 2023***) can make overconfident predictions. In contrast, trustworthy, well-calibrated models should exhibit high uncertainty when predictions are likely to be inaccurate and low uncertainty when they are likely to be accurate. In neuroscientific applications, the ability to assess uncertainty can be particularly important, for example in tasks where wrong predictions can have serious consequences, such as brain-computer interfaces and real-time decoding for an actuator.

Classical VAEs provide principled access to the predicted uncertainty over the inferred latent states, often in the form of the variance of a Gaussian approximate posterior distribution (***Rezende et al., 2014***; ***Kingma and Welling, 2014***). However, this aspect is sometimes treated as a convenience for robust training rather than as a meaningful quantity with respect to the system under investigation. In this work, we have demonstrated the effect of increased latent uncertainty when dealing with partially observed inputs, highlighting the need for accurate modeling of latent uncertainty.

Further, the observation noise process can be modeled explicitly in VAEs allowing the combination of different data types (***Nazábal et al., 2020***; ***Brenner et al., 2024***): e.g., Poisson noise for spike counts and Gaussian noise for behavioral trajectories. Using a Gaussian negative log-likelihood loss allows to estimate observation noise for each data channel separately, a desirable property in scientific measurements, yet challenging to accomplish (***Rybkin et al., 2021***; ***Seitzer et al., 2022***).

To assess overall calibration in VAEs, which reflects both the posterior uncertainty and observation noise, we introduced a version of simulation-based calibration (***Talts et al., 2018***; ***Cook et al., 2006***), which allows for sample-based uncertainty evaluation in the absence of tractable ground truth distributions. Assessing calibration for discrete data poses additional challenges—in particular in a low count regime where it is not possible to obtain reliable confidence intervals—and remains an avenue for future investigation (***Wei and Held, 2014***).

## Limitations

Our masked VAE approach allows for calibrated predictions on a variety of conditioning tasks, but has some limitations: First, our approach relies on a small number of specified and *structured* conditioning masks, rather than considering all possible combinatorial (2^*D*^) masking conditions, where *D* is the data dimension. For many problems in neuroscience, this suffices since the conditional distributions of interest are usually few and well-specified, such as behavior given neural data. To tackle the problem of capturing *all* conditionals, it would be an empirical question of how big models and datasets would need to be to effectively generalize to this combinatorial space. Second, similar to other deep-learning-based methods, our approach struggles with capturing interactions on long and varying timescales. Here, we only investigated cross-modality interactions occuring on similar timescales (reach movements) and fixed the length of the time segments during training (maximum of 150 time steps). Thus, behavior or neural activity preceding this segment cannot influence subsequent predictions. Integrating transformer-based approaches (***Vaswani et al., 2017***; ***Jaegle et al., 2022***) might be useful for capturing such interactions over varying timescales (***Ye and Pandarinath, 2021***; ***Ye et al., 2023***; ***Azabou et al., 2023***; ***Antoniades et al., 2024***).

Third, our approach inherits common issues from VAEs, for example, the lack of a principled way to choose the dimensionality of the VAE latent space, rendering hyperparameter tuning potentially costly. If the dimensionality of the latent space is too large, the VAE might fail to exploit correlations within the data and use separate latent variables for the specified conditional distributions, potentially degrading the quality of conditional generation. On the other hand, if the dimensionality is too small, the VAE might not be able to accurately model the data. Thus, for the monkey reach task, we introduced a sparsity-inducing prior (***Ainsworth et al., 2018***) that mitigates this issue by automatically reducing the latent dimensionality if latents are not used by the decoder network.

Lastly, while samples from our masked VAE are well-calibrated in most cases and often close to the ground-truth neural and behavioral trajectories, the sampling quality of VAEs is known to be limited even for simpler and fully observed datasets. Recent generative models such as Denoising Diffusion Probabilistic Models (***Ho et al., 2020***; ***Rombach et al., 2022***), Normalizing Flows (***Rezende and Mohamed, 2015***), and Generative Adversarial Networks (***Goodfellow et al., 2014***) can produce samples of higher quality, but they lack the main feature of VAEs that makes them especially relevant in neuroscience: inference of low-dimensional latent states. Combining our approach with other such generative models (e.g., ***Zhou and Wei*** (***2020***); ***Bashiri et al.*** (***2021***)) could be an interesting future avenue to improve sampling quality while preserving the possibility of performing latent inference.

## Conclusion

We present a method that addresses two common goals in neuroscience: inferring low-dimensional representations and unveiling dependencies in simultaneously recorded modalities by modeling their conditional distributions. Our approach will allow for scaling encoding and decoding analyses in neuroscience to today’s high-dimensional multi-modal datasets. Furthermore, this work highlights a crucial aspect of analyzing neural and behavioral data: the importance of uncertainty estimates.

## Materials and Methods

Here, we adapt variational autoencoders (***Kingma and Welling, 2014***; ***Rezende et al., 2014***) to address two goals simultaneously: First, to infer low-dimensional representations underlying multi-modal neural and behavioral time-series data and, second, to model their conditional distributions. Modeling conditional distributions is ubiquitous in neuroscience, and since neuroscientific data are typically variable even in controlled experiments, relations between modalities may also be variable. Therefore, we focus on probabilistic rather than deterministic approaches to characterize such conditional distributions. We reformulate the estimation of conditional distributions in VAEs in a more general way: modeling the distribution of an unobserved subset of the data given an observed subset *p* ( unobserved observed ) similar to (***Collier et al., 2020***; ***Nazábal et al., 2020***). To target such distributions with a VAE, we modify the training scheme and loss of classical VAEs. We validate our approach on a tractable example and two neuroscientific time-series datasets: walking behavior of the fly and a continuous reach task in monkeys. We introduce calibration metrics to evaluate the models’ uncertainty estimates in the context of scientific data, i.e., without access to ground-truth uncertainties.

### Background on Variational Autoencoders

Variational Autoencoders (VAEs) are probabilistic models capable of capturing complex multi-modal data distributions *p*(**x**). The assumption underlying VAEs is that all variations in the data distributions can be captured (up to observation/measurement noise) by the variations of corresponding unobserved latent variables *z*. VAEs learn stochastic mappings between the observed data space and the unobserved or latent (*z*-space). Both mappings from the data to latent distributions and vice versa are typically parameterized through flexible neural networks. The generative model is described by the joint distribution of data and latent variables, which factorizes into

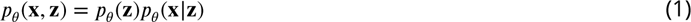

parameterized by *θ* where *p*(**z**) is the prior over the latent space. The prior is usually chosen to be a simple distribution such as a standard Gaussian, and *p*_*θ*_ (**x ∣ z**) is the probabilistic decoder. The inference or encoder model *q*_*ϕ*_ (**z ∣ x**), parameterized by *ϕ*, which infers the latent distribution from data, is an approximation of the true, intractable posterior *p* (**z ∣ x**) (***Kingma and Welling, 2014***; ***Rezende et al., 2014***; ***Kingma and Welling, 2019***). VAEs are trained by

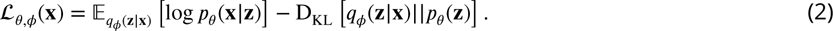

The first term assesses how well the predicted distribution matches the original data and is often referred to as the reconstruction loss. The second term is the Kullback-Leibler divergence D_KL_ between the approximate posterior *q*_*ϕ*_(**z**|**x**) and the latent prior *p*(**z**), which regularizes the learned latent space. Maximizing the ELBO *L*_*θ*,*ϕ*_(**x**) with respect to the parameters *θ* and *ϕ* leads to a better generative model and increases the similarity between the approximate and the true (intractable) posterior. All parameters *θ* and *ϕ* can be optimized jointly using stochastic gradient descent. Once trained successfully, one can sample from the prior and consecutively from the stochastic decoder output to obtain a new sample *x*^_pred_ from the learned data distribution. Alternatively, one can sample from the approximate posterior of a previously unseen test datum *x*_test_ to obtain reconstructions that closely resemble *x*_test_.

VAEs have been successfully applied to various types of potentially heterogeneous data (continuous, discrete, ordinal, etc.) (***Nazábal et al., 2020***) and have been extended to time series (***Gregor et al., 2015***; ***Chung et al., 2016***; ***Girin et al., 2021***), paving the way for applications to neuroscientific time series (***Sussillo et al., 2016***; ***Pandarinath et al., 2018***; ***Luxem et al., 2022***; ***Brenner et al., 2024***).

### Capturing arbitrary conditional distributions with VAEs

To model flexible conditional distributions with VAEs, we modified the training scheme of classical VAEs similar to (***Nazábal et al., 2020***; ***Collier et al., 2020***). While training the VAE, we randomly mask out subsets of the data corresponding to the desired conditional distributions and compute the loss on the remaining data. Prior to training, for each conditional distribution of interest, we specify a conditioning mask *m* together with a mask probability *p*_*m*_ (***Figure 1c***, left). During training, the conditioning masks are sampled independently for each data point according to the mask probabilities. Concretely, the masking is performed by replacing the data with their respective mean values. Other replacement values, such as zeros for spiking (count) data, are also possible. We calculate the reconstruction loss *L_recon_* solely on observed, that is, non-masked data. Sometimes, to facilitate learning that some data has been masked out, we additionally provide the encoder network with a binary mask consisting of 0s for unobserved and 1s for observed data points (see networks).

Through this training procedure, the masked VAE simultaneously optimizes the ELBO over all different conditional distributions, that is

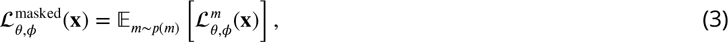

where *p*(*m*) is the previously specified probability distribution over all conditioning masks, including the fully observed case, where no data is masked out. As noted above, the mask *m* is applied to both the data and the corresponding part of the reconstruction loss. This training procedure promotes the learning of different encoder networks that share parameters, allowing us to target different approximate posterior distributions given different conditioning masks.

From an implementation perspective, the conditioning masks can be passed to the encoder network in various ways. They can, for example, be concatenated or added directly to the input, but also at later stages in the network, possibly after transformations with a (non-linear) embedding. In contrast to (***Collier et al., 2020***), we explicitly do not pass the conditioning masks to the decoder since all uncertainty and mean shifts induced by masking should be reflected in the latent representation.

### Modeling observation noise with VAEs

It is important to ensure that VAEs correctly capture the uncertainties in the (conditional) data distributions. In a VAE, there are two sources of uncertainty - the inferred posterior uncertainty and the observation or measurement noise. The latter source of uncertainty is often ignored, which is reflected in the common choice of the mean squared error (MSE) as the reconstruction loss. The MSE only evaluates the quality of the mean prediction and ignores the stochastic nature of the VAE decoder. If we instead want to correctly capture the observation noise, it is necessary to learn it explicitly. Assuming that the observation noise follows a Gaussian distribution, we use the Gaussian negative log-likelihood (GNLL) as our reconstruction loss. The GNLL for an observation *x* given a model prediction of the Gaussian mean *μ* and standard deviation σ is given by

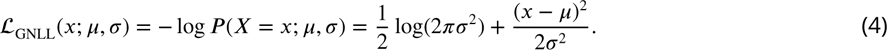

Note that the MSE is a special case of the GNLL where the standard deviation is set to 1. As noted above, this usually leads to samples from the model that are not calibrated in the statistical sense. Optimization, however, is more challenging when using the GNLL and might require additional adjustments (***Rybkin et al., 2021***; ***Seitzer et al., 2022***).

### Datasets and Data Preprocessing

#### Linear Gaussian Latent Variable Model

We simulated a dataset based on a Gaussian Latent Variable Model (GLVM) with one latent variable *z*, where *z* ∽ *N*(0, 1), and 20 data dimensions **x**, where **x ∽ *N* (C*z* + d, ⋀)** (***Figure 2a***). To demonstrate the difference between noisy and more precise, less noisy, variables in a setup that accounts for uncertainty, noise levels for all data dimensions differ. For each data dimension *i*, σ_*i*_ is drawn from a log-normal distribution with *μ*_*LN*_ = log(0.7) and σ_*LN*_ = 0.5. *C*_*i*_s are drawn from a normal distribution with *μ*_*N*_ = 1.1 and σ_*N*_^2^ = 0.1. Additionally, the sign for a given *C*_*i*_ is flipped with probability 0.5. All offsets **d** are set to 1. We use 9000 samples from this model for training and 1000 each for validation and testing. This fully Gaussian setup allows for the analytical computation of both conditional *p*(**x**^unobs^|**x**^obs^) and posterior *p*(*z*|**x**^obs^) distributions (see Appendix), which we compare to the distributions learned by our model.

#### Fly walking behavior

To collect the data on fly walking behavior, we placed flies, *Drosophila melanogaster*, in an acrylic arena that constrained them to move in a 2D plane. Thus, flying is not part of the otherwise rich repertoire of observed behaviors, which includes both forward and backward walking, grooming, resting, etc. The flies were part of a genetic screen (not wild-type) but were examined during behavior capture and were morphologically and behaviorally indistinguishable from wild-type flies. We placed three female flies in one arena simultaneously and filmed them from below, three times for three seconds (frame rate of 80*Hz*). This procedure was repeated about 10000 times, resulting in 28059 time series with 234 time points each. To extract each fly from the video separately, we tracked the centroid of each fly using Tracktor (***Sridhar et al., 2019***), cropped out the flies in each frame, and aligned them to point in the upward direction. We then tracked 32 body parts (four joints per leg, as well as head features, thorax, abdomen, and wings), each with x- and y-directions using DeepLabCut (***Mathis et al., 2018***), resulting in time series with 64 feature dimensions. We then smoothed the extracted time series using a Savitzky-Golay-Filter (***Savitzky and Golay, 1964***) with a polynomial order of two and a window length of seven. The smoothed trajectories were then cut into sequences of length 48 with buffers of length 9 between each sequence to avoid information leakage. 95% of the data was used for training, while the remaining data was used for validation and testing (2806 sequences each). Potential information leakage due to autocorrelation between training and test/validation sets is further reduced by choosing the last sequences for testing/validation instead of an interleaved approach, which can often cause information leakage in time-series models. Prior to passing the time series to the network, we standardize each feature dimension across the 48 time steps.

#### Continuous reach task in monkeys

The neural and behavioral dataset, made publicly available by ***O’Doherty et al.*** (***2017***), was recorded from two monkeys (rhesus macaque) performing self-paced continuous reaches, i.e., without gaps or pre-movement delay intervals. Targets were arranged in an 8 by 8 grid, and a new target was presented when the previous target was reached. Neural recordings were taken from the cortical hemisphere contralateral to the arm performing the reach movements. ***O’Doherty et al.*** (***2017***) provide the neural data after spike sorting in the shape channels vs. spike times. We focus only on one session, ‘loco_20170213_02‘, which contains neural activity from both primary motor (M1) and (S1) activity, as well as the cursor, target, and finger positions. Here, we take only the neural activity from M1 and the cursor positions (*x*,*y* direction) as the behavioral correlate. We filter out channels with firing rates below 0.5 Hz analogous to (***Makin et al., 2018***), resulting in 213 remaining M1 units. We convert spike times into spike counts in bins of 64 ms (15.625 Hz). We down-sample the cursor position by querying it at fewer time points consistent with the reduced sampling rate used for binning the spikes (15.625 Hz instead of 250 Hz). We do not introduce a delay between neural activity and behavior as done, e.g., in (***Schimel et al., 2021***; ***Jensen et al., 2021***), we rather let the model identify which aspects of the respective other time series to consider for its predictions. We use the first 70% of the data for training (approx. 28 minutes recording time), the following 10% for testing (approx. 4 minutes), and the remaining 20% for validation (approx. 8 minutes). We standardize the behavioral train, test, and validation time series with respect to the overall mean and standard deviation of the training set for both reach directions. During training, we introduce ‘pseudo-trials’ with 150 time steps each, that start at randomly sampled time points.

### Network Architectures and Optimization

Model details: GLVM

**Training scheme and masks:** Prior to training, we select three randomly sampled masks (10 out of 20 dimensions are masked) to test if the masked approach can capture the true posterior and conditional distributions. Since we chose different loading factors *C*_*i*_ and noise levels σ_*i*_ for each dimension, the corresponding posterior mean and variance, and thus also the conditional distributions, differ between conditions. During training, we uniformly sampled the four conditioning masks (all observed and mask 1-3) and used the Adam optimizer (***Kingma and Ba, 2015***) to train our model.

**Architecture:** The encoder network consists of a simple linear embedding for the mask, which is passed through a multilayer perceptron together with the 20-dimensional data vector to parameterize the one-dimensional posterior mean and log variance. To focus on posterior inference under different masking conditions, we set the decoder to be the true generative model. Note, however, that this GLVM example is identifiable, i.e., all parameters of the generative model (*C*_*i*_, *d*_*i*_, σ_*i*_) can be learned using a VAE, which we confirmed even in our masked training scheme. Nevertheless, fixing the decoder is beneficial in this case since the posterior is only identifiable up to a rotation in the latent space (*μ*_*z*_, σ_*z*_), which in the one-dimensional setting corresponds to a flipped sign.

**Loss:** For this well-specified, identifiable Gaussian example, the regular masked GNLL was used together with a standard Gaussian prior in the latent space.

### Model details: Fly walking behavior

**Training scheme and masks:** To investigate low-dimensional representations of fly walking behavior, we built a sequential VAE and specified masks for the body keypoints most relevant to walking. Analogous to the GLVM case, we adapted the masked training scheme for the time-series case to allow for modeling the conditional distribution over a subset of the fly body keypoints, given the remaining keypoints. Specifically, we mask the hind claw, hind tibia-tarsal joint, mid tibia-tarsal joint, and mid claw of the left side. The entire time segment of masked keypoints is replaced with the mean value across this segment. We assign a probability of 50% to the all-observed and the leg-masking condition. We again use the Adam optimizer (***Kingma and Ba, 2015***) with a learning rate of 0.0005 for training our model.

**Architecture:** The VAE for fly walking behavior consists of an encoder and a decoder network that are both trainable neural networks. The encoder network consists of two sets of 1D convolutional layers, each followed by batch normalization and ELU activation. We then apply temporal convolutions that compress the data in the temporal dimension before passing it to a bidirectional RNN (***Cho et al., 2014***) for temporal context. Thus, the encoder network is non-causal in time. The RNN output is then passed through a multi-layer perceptron to parameterize the posterior mean and log variance. This results in a latent space with spatial (*N*_*z*_) and temporal (*T*_*z*_) dimensions smaller than the 64 features and 48 time-steps of the data (for our choice of parameters *N*_*z*_ = 18 and *T*_*z*_ = 13, i.e., the size of the latent space is less than 8% of the original data). Unlike in the GVLM case, we do not pass the mask to the encoder network, since it does not improve the conditional modeling. After sampling from the approximate posterior, the decoder network expands the time dimension of **z** using transposed convolutions, followed by dimensionality expansion to parameterize the Gaussian mean and observation noise variance. The latter is constrained to be positive by a softplus function to ensure well-defined variances.

Loss:

For the continuous behavioral data, we again use GNLL (eq. 4), which is computed per feature and timepoint. The prior distribution in the latent space is standard Gaussian.

### Model details: Neural and behavioral data from a monkey reach task

**Training scheme and masks:** The sequential VAE for the monkey reach task jointly models time series of high-dimensional neural spike-count data and continuous cursor positions. We specify the masks required for neuroscientific encoding and decoding: either all neural activity is masked out (spike counts set to zero), or all behavioral traces are masked and set to their respective mean values. Following ***Ainsworth et al.*** (***2018***), we introduced a sparsity-inducing prior that sets latent contributions to zero if they are not used by the model. We used the AdamW optimizer (***Loshchilov and Hutter, 2019***) with a learning rate of 0.001 and weight decay of 0.2) to train our encoder and decoder networks. For parameters related to sparsity-induction, we follow (***Ainsworth et al., 2018***) and use Stochastic Gradient Descent (SGD) with zero momentum. Here, we show results that are trained on only one session, but the architecture allows training on data from multiple sessions using session-specific input and output mappings.

**Architecture:** First, we expand the data using a session-specific linear mapping. Similar to the sequential VAE for the fly data, the encoder network then performs a non-linear dimensionality reduction followed by a bidirectional RNN to parameterize the latent posterior mean and log-variance. Here, we do not consider compression in time, and each latent time-point corresponds to a time-point in dataspace. The decoder also has an RNN and uses further multi-layer-perceptrons to map the latent samples back into data space. For the continuous behavioral data, the decoder again predicts the Gaussian mean and observation noise variance. For the discrete spike data, however, the decoder only models the underlying firing rates.

**Loss:** This discrepancy arises from the different distributions used to model the respective data modalities. While behavior is continuous and thus appropriately modeled with a Gaussian, discrete spike counts are best modeled with a Poisson distribution. Consequently, the GNLL is replaced by the negative Poisson log-likelihood, which, for an observed spike count *x* and a rate parameter *λ*, is defined as:

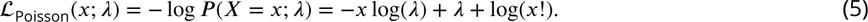

For this task, we also refined the behavioral GNLL loss by incorporating the concept of *β*-NLL introduced by ***Seitzer et al.*** (***2022***) by weighting each data point’s contribution to the loss based on the *β*-NLL-exponentiated variance estimate σ_*i*_^2*βNLL*^. Effectively, this small loss modification prevents a potential issue when learning the observation noise, namely that poorly fitted variables are assigned high variance, which, since it appears in the denominator in eq. 4, leads to smaller gradients and hence less incentives for the network to improve its fit. In this application, a NLL-beta of 0.3 worked well. Before calculating the overall gradients, we sum up the behavioral and neural contributions to the reconstruction loss. Note that when using heterogeneous noise models the scales of the loss contributions can vastly differ. Hence, depending on the application and downstream tasks, it can be beneficial or even necessary to introduce weighting factors to balance out the losses and corresponding gradients (***Javaloy et al., 2022***). To improve stability and prevent over-fitting, we additionally regularize the session input and output weight matrices. For the sparsity-inducing prior in the latent space, we apply a version of Lasso regularization that encourages sparsity in the weight matrix that transforms the *z*-samples before they are passed through the decoder (see ***Ainsworth et al.*** (***2018***) for details).

### Analysis and Metrics

#### Calibration metrics: Evaluating uncertainties in variational autoencoders

To investigate the statistical calibration in VAEs, we perform a version of simulation-based calibration (***Talts et al., 2018***; ***Cook et al., 2006***), which is associated to frequentist coverage tests (***Wei et al., 2023***). Here, we focus on the calibration of the predictive distribution in data space. For each test datum *x*_test_, we sample *n*_*z*_-times from the approximate posterior, pass the sampled *z* through the decoder and sample from the observation noise model *n*_*obs*_-times. This sampling procedure results in a sampling distribution in data space that reflects both posterior uncertainty and observation noise.

For continuous data, we then compute confidence intervals corresponding to the *n*th percentile. For a statistically well-calibrated model, *n*% of the ground truth data should lie in this interval. When plotting the different percentiles against the proportion of data points falling in the corresponding interval, well-calibrated predictions lie on the diagonal. Here, we evaluate this for the 60th, 80th, 90th, and 95th percentiles, which roughly correspond to one to three standard deviations. However, other evaluations, including percentile bins from 0 to 100, are also common (see e.g. ***Wei et al.*** (***2023***)). If a model is overconfident, the corresponding values fall in the lower triangle below the diagonal. In the underconfident regime, i.e., if the average predictions are accurate, but the estimated uncertainty is too high, the values fall in the region above the diagonal. In all three datasets, we evaluate the calibration of masked continuous variables (GLVM, fly walking behavior, and behavior in the monkey reach task). The test set sizes, as well as the computational cost of estimating confidence intervals, differ between models. Therefore, we chose different *n*_*z*_ and *n*_*obs*_ to compute the confidence intervals but kept both values the same when evaluating the naive model to ensure a fair comparison.

For count data, we compute cumulative distribution functions (CDFs) of the spike counts to assess the calibration of the predicted firing rate. This is because most bin counts are either 0 or 1, making it impractical to construct confidence intervals (***Wei and Held, 2014***). More specifically, to obtain informative CDFs, we aggregate five neighboring bin counts. Then, for each of the *n*_*z*_ ⋅ *n*_*obs*_ predicted rate time series, we sample spikes and compute the CDF over all 40 aggregated bins. Finally, we plot the obtained CDFs against the analogously aggregated CDF of the ground truth spike train. If the rate predictions are well calibrated, their resulting CDFs closely match the ground truth CDF and lie on the diagonal.

### Code and data availability

Datasets and code for this study will be made available at https://github.com/mackelab/neuro-behavior-conditioning upon publication.

## Acknowledgements

This work was supported by the German Research Foundation (DFG) through Germany’s Excellence Strategy (EXC-Number 2064/1, PN 390727645) and SFB1233 (PN 276693517), SFB 1089 (PN 227953431), the German Federal Ministry of Education and Research (Tübingen AI Center, FKZ: 01IS18039; the Human Frontier Science Program (HFSP), and the European Union (ERC, DeepCoMechTome, 101089288). DM acknowledges a Marie Curie EuroTech postdoctoral fellowship, a Swiss Government Excellence Postdoctoral Scholarship (2018.0483) and funding from the European Union’s Horizon 2020 research and innovation program under the Marie Skłodowska-Curie grant agreement no. 754462. VL-R acknowledges support from the Mexican National Council for Science and Technology, CONACYT, under the grant number 709993. PR acknowledges support from an SNSF Project grant (no. 175667) and an SNSF Eccellenza grant (no. 181239). AS and JV are members of the International Max Planck Research School for Intelligent Systems (IMPRS-IS). We would like to thank Paul Fischer for data management support for the fly dataset, Lisa Haxel for feedback on the manuscript, and all Mackelab members for discussions and feedback throughout the project.

## Supplementary Figures

**Figure S1.**
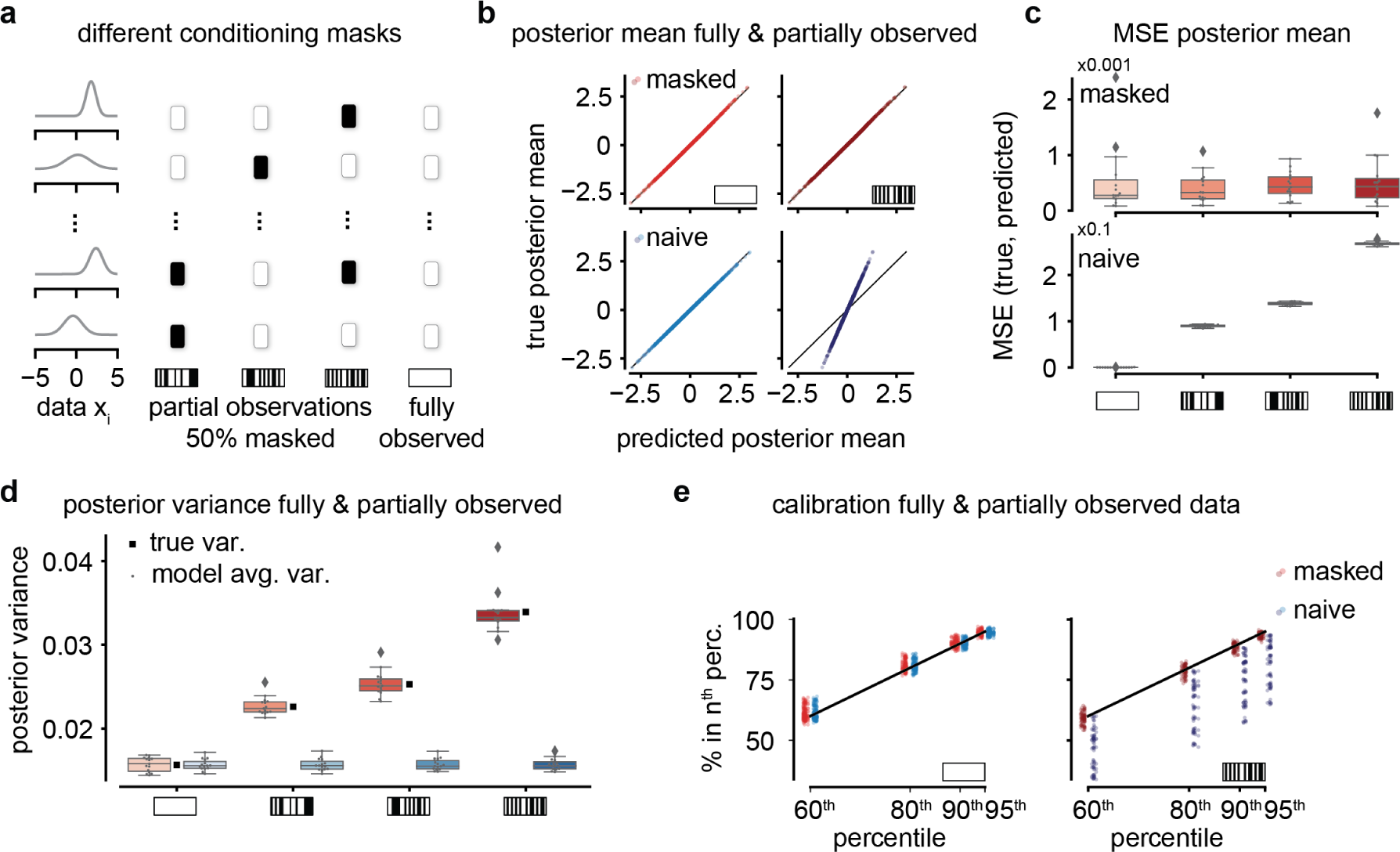
Posterior inference for different masking conditions a We test three different masking conditions where half of the input data dimensions are masked in addition to the fully observed case. b Masked VAEs correctly capture the posterior mean for partially and fully observed conditions, while the naive VAE can only perform correct inference in the fully observed condition. c The mean squared error (MSE) between the true and predicted posterior mean is low for all conditions for masked but not naive VAEs. Note the different orders of magnitude. d Masked VAEs appropriately adjust the posterior variance for all conditions, while masked VAEs always wrongly predict the variance level of the fully observed condition (white rectangle). e Both naive and masked VAEs are well calibrated in the fully observed condition (left). Yet, only masked VAEs are calibrated when modeling conditional distributions (right).

**Figure S2.**
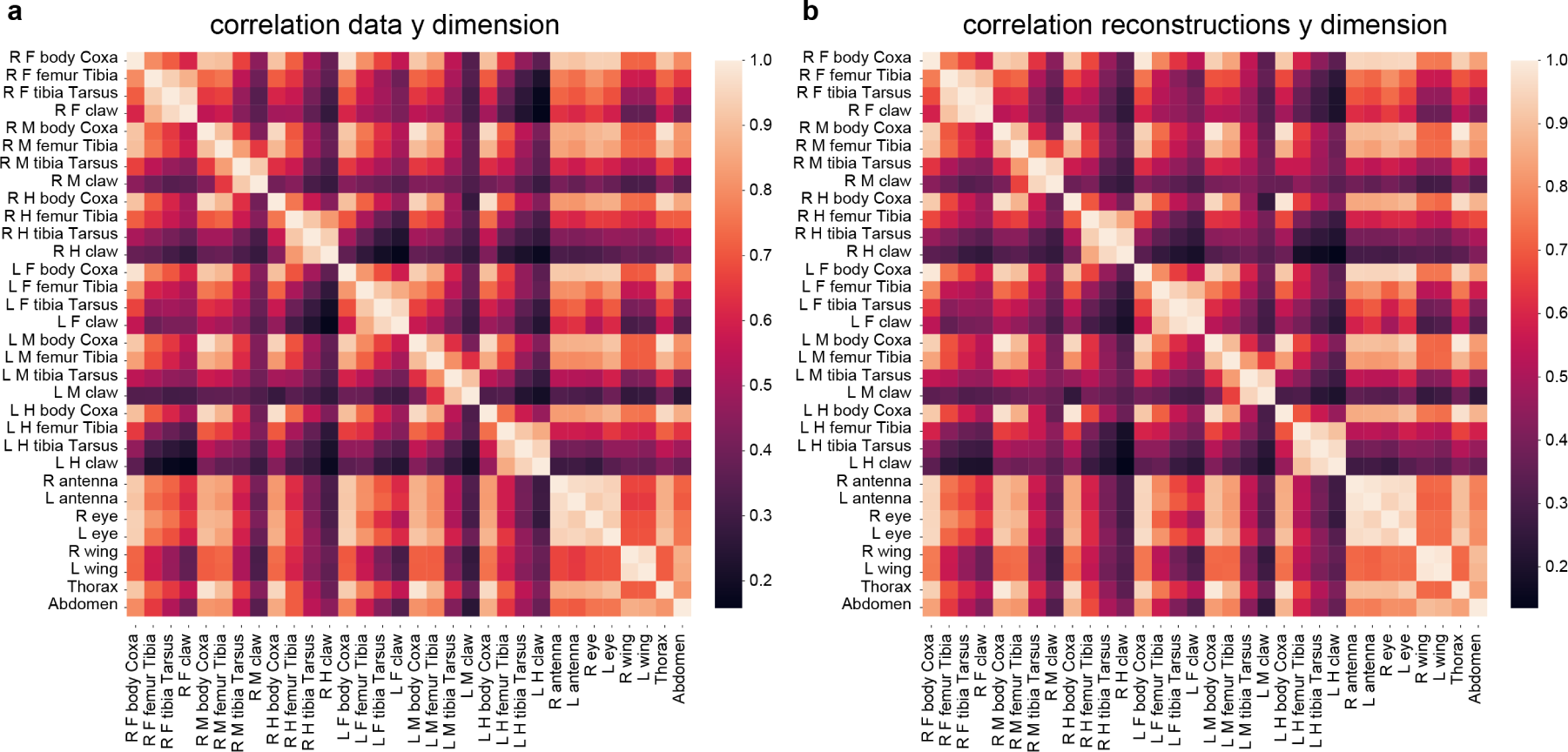
The VAE correctly captures the correlation structure between the fly keypoints. a Correlation matrix of the y-dimension of the test set. b Correlation matrix of the y-dimension of the corresponding VAE reconstructions (mean)

**Figure S3.**
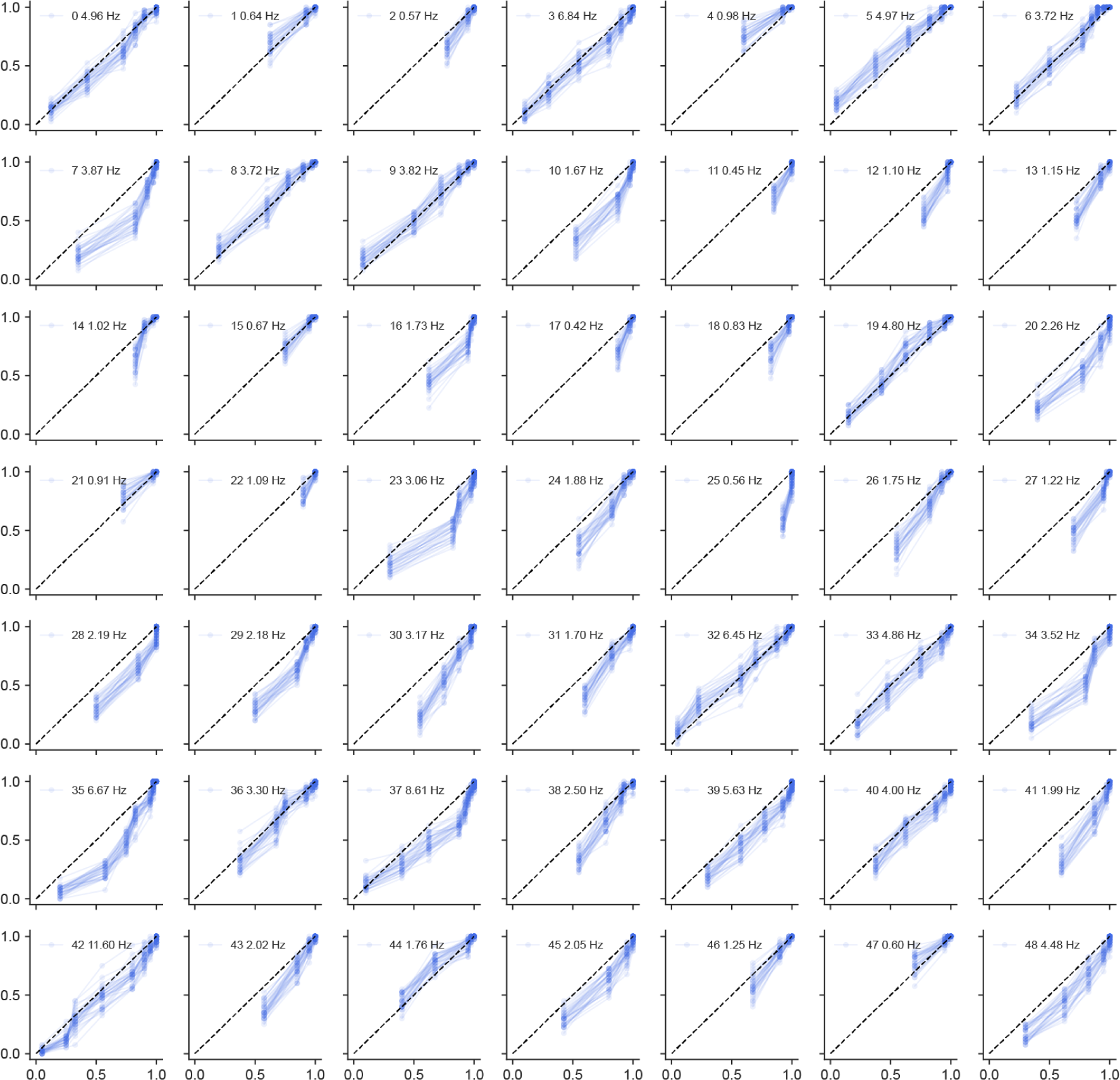
Additional cumulative distribution functions of units from monkey primary motor cortex Subset of 49 units. Sample CDFs of the encoding predictions of the naive VAE.

**Figure S4.**
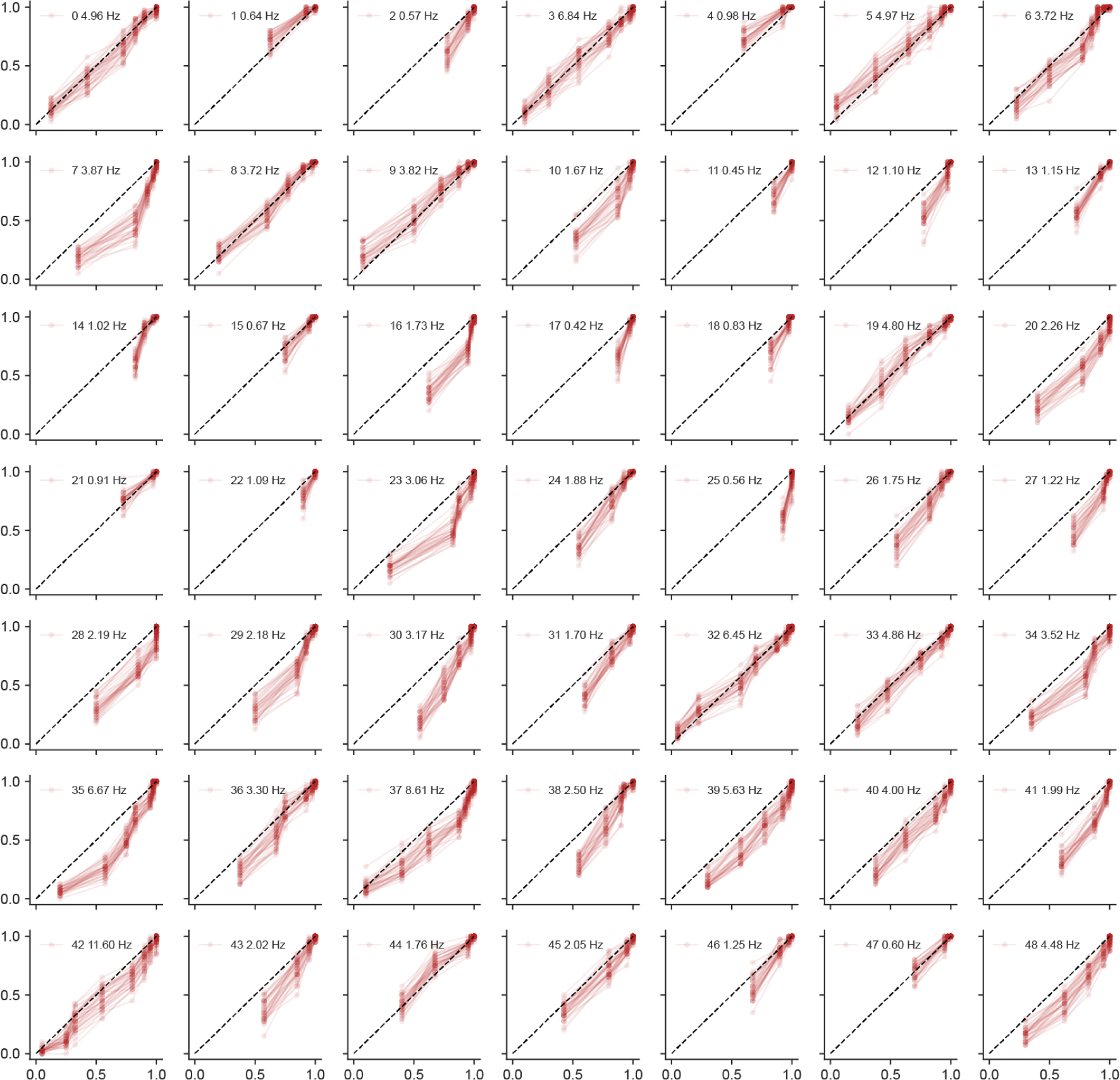
Additional cumulative distribution functions of units from monkey primary motor cortex Subset of 49 units. Sample CDFs of the encoding predictions of the masked VAE.

**Figure S5.**
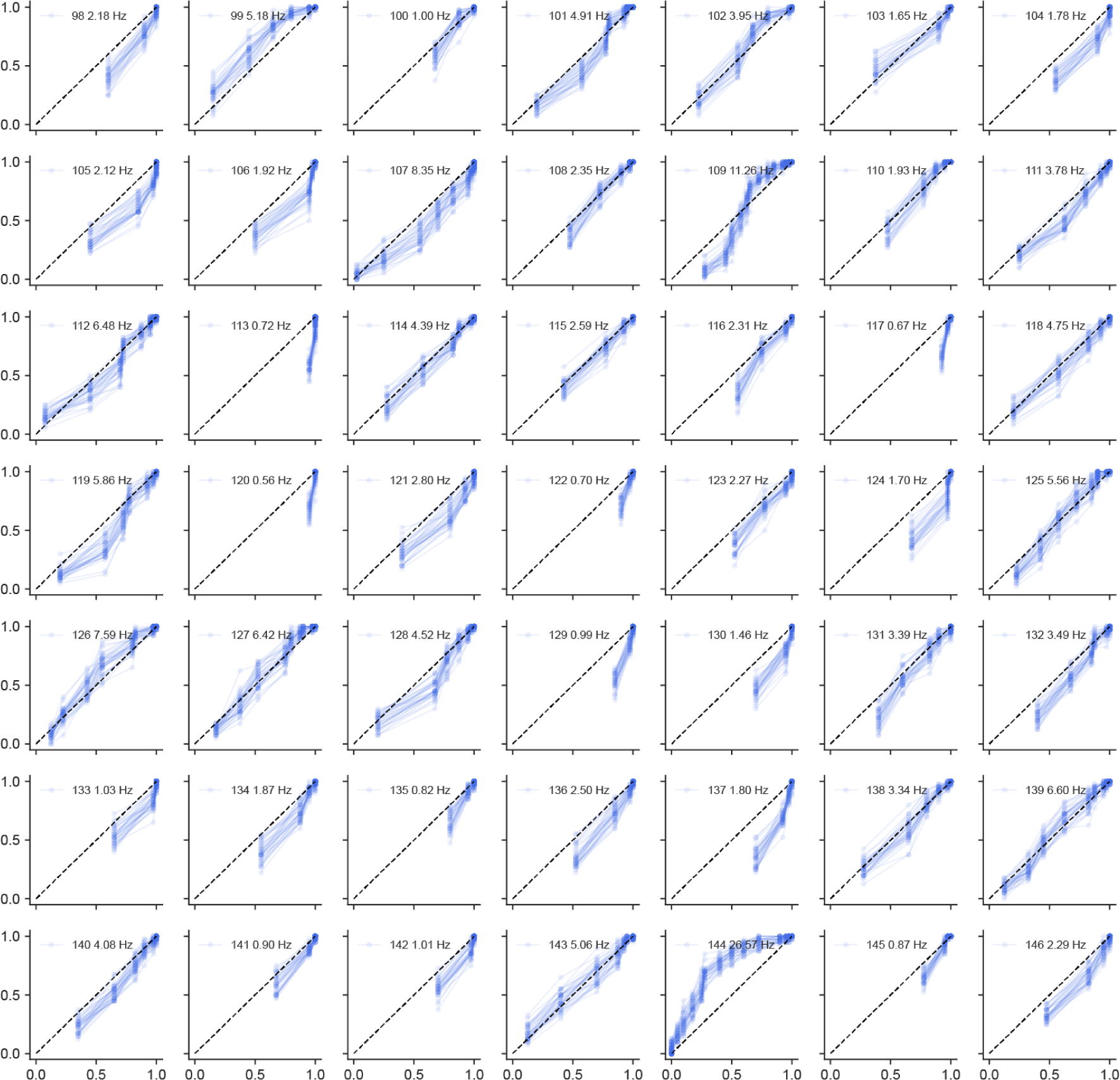
Additional cumulative distribution functions of units from monkey primary motor cortex Subset of 49 units. Sample CDFs of the encoding predictions of the naive VAE.

**Figure S6.**
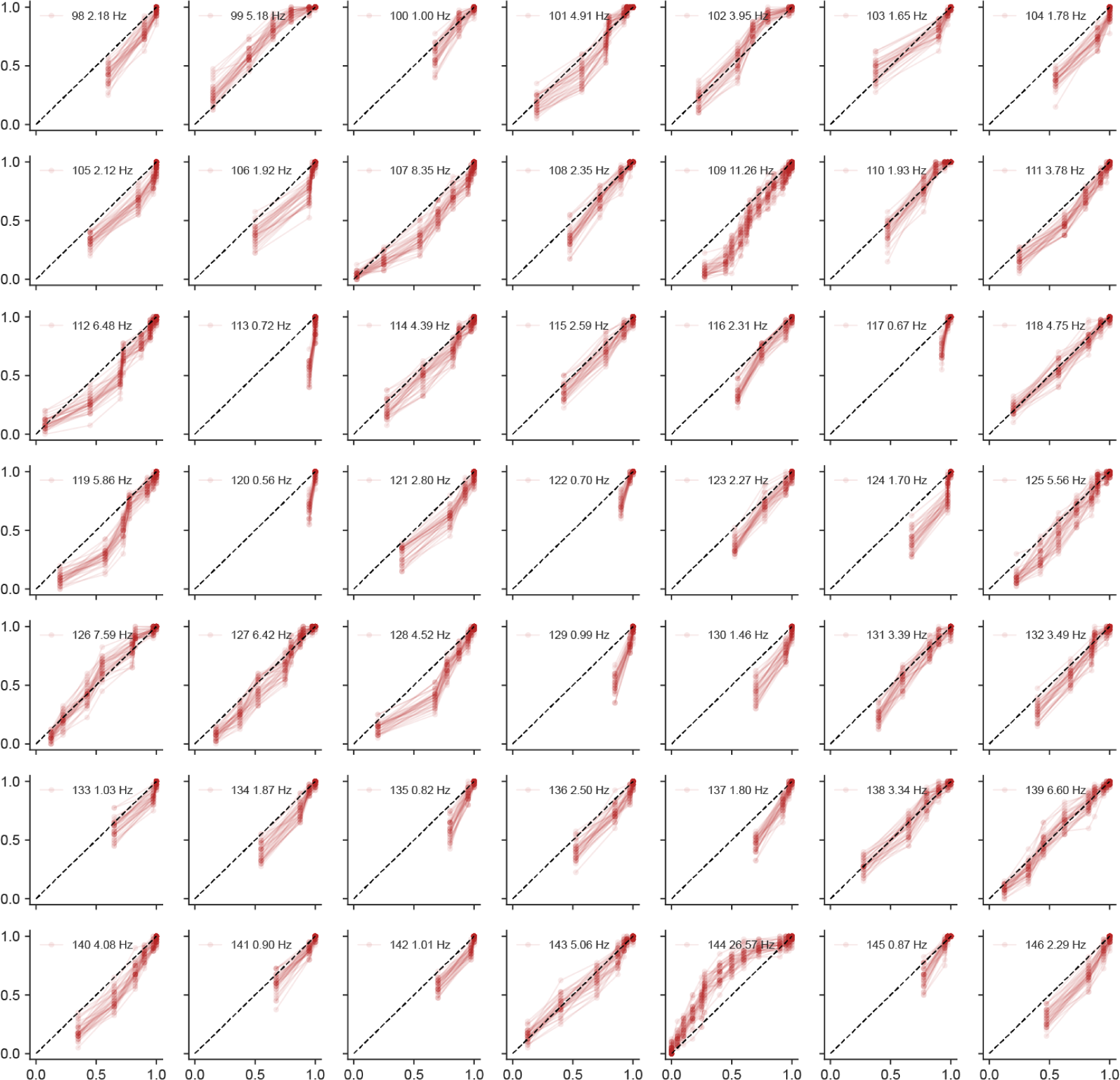
Additional cumulative distribution functions of units from monkey primary motor cortex Subset of 49 units. Sample CDFs of the encoding predictions of the masked VAE.

## Supplementary Dataset Information and Hyperparameters

**Table S1.** Summary of relevant hyperparameters and dataset information for the GLVM VAE training.

| Hyperparameter | Value |
| --- | --- |
| Learning Rate | 0.001 |
| Beta | 1.0 |
| Warmup Range | 30 epochs |
| Beta Step | 1/30 |
| Epochs | 1500 |
| Fraction fully observed | 0.25 |
| Latent Size | 1 |
| Training Batch Size | 1000 |
| Validation/Test Batch Size | 1000 |
| Dataset dimensions (samples, dimensions) | (10000, 20) |
| Trainable Parameters | 7442 |

**Table S2.** Summary of relevant hyperparameters and dataset information for the fly VAE training.

| Hyperparameter | Value |
| --- | --- |
| Learning Rate | 0.0005 |
| Beta | 1.0 |
| Warmup Range | epochs 3-6, beta=0 before |
| Beta Step | 0.25 |
| Epochs | 700 |
| Epoch when masking starts | 250 |
| Fraction fully observed | 0.5 |
| Latent Size | 18 |
| Training Batch Size | 256 |
| Validation/Test Batch Size | 512 |
| Dataset dimensions (samples, sequence length, key points) | (28059, 234, 64) |
| Sequence length cut for training | 48 |
| Trainable Parameters | 39426 |

**Table S3.** Summary of relevant hyperparameters and dataset information for the monkey reach VAE training.

| Hyperparameter | Value |
| --- | --- |
| Learning Rate | 0.001 |
| Learning Rate sparsity | 0.005 |
| Beta | 1.0 |
| Warmup Range | 50 iterations |
| Beta Step | 1/50 |
| NLL Beta | 0.3 |
| Iterations | 25000 |
| Iteration when masking starts | 5000 |
| Fraction fully observed | 0.5 |
| Max. Latent Size | 40 |
| Training Batch Size | 256 |
| Validation/Test Size | 512 |
| Dataset dimensions | total time: 26439, units: 213, behavior: 2 |
| Sequence length cut for training | 150 |
| Trainable Parameters | 379177 |

## Supplementary Information Gaussian Latent Variable Model

Here we consider a linear Gaussian latent variable model with a one-dimensional latent space and n-dimensional observations *x*.

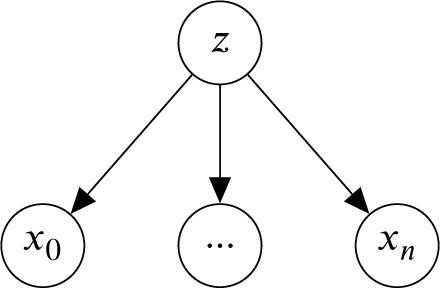

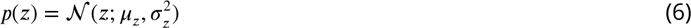

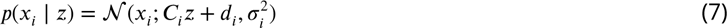

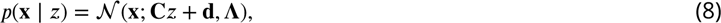

where ∧) = diag(σ_*i*_^2^).

### Joint distribution of latent and observed variables

For a GLVM as outlined above the joint distribution of latent and observed variables can be written as

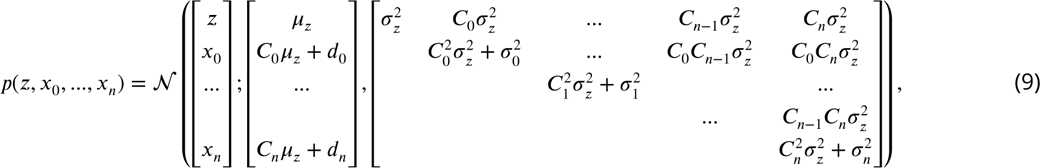

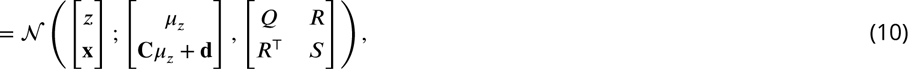

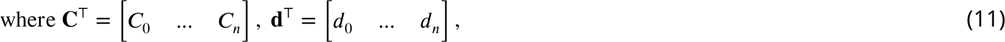

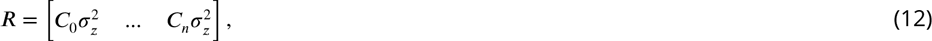

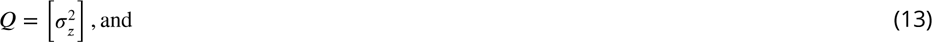

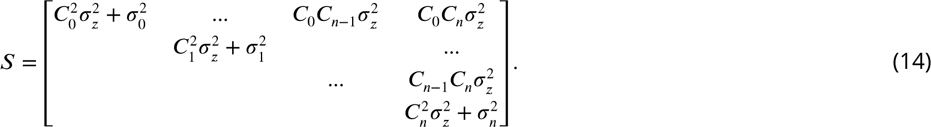

Marginalization in the fully Gaussian case corresponds to ignoring the rows and columns of the mean and covariance matrices of the joint distribution that correspond to the variable we aim to marginalize out. This means that the covariance of e.g. the joint of only the data dimensions *p*(*x*_0_, …, *x*_*n*_) is simply the submatrix that leaves out the first row and column of (9). Corresponding adjustments to the matrices when marginalizing out certain variables are denoted as 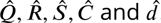.

### Posterior distribution - inferring the latent distribution given observations

In order to compute the posterior of *p*(*z* ∣ **x**), we can apply the conditioning rule for Gaussians and obtain

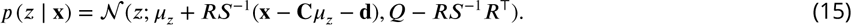

If some variables are unobserved, we can still perform exact inference of the posterior distribution *p*(*z* ∣ **x**^obs^). We obtain the analytical result by first marginalizing out the respective unobserved variables in the joint distribution in Eq. 9 before performing the conditioning step as in Eq. 15.

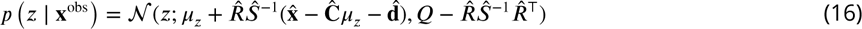

where the hat indicates altered matrices where respective entries corresponding to unobserved values have been dropped.

Hence, for each masking pattern, a different matrix inversion has to be carried out. Also, see the discussion of exact inference in the factor analysis model in ***Williams et al.*** (***2019***). It is worth noting here that the posterior variance does not depend on the exact *x* values. Intuitively, the posterior variance (uncertainty) will increase if some of the inputs are unobserved.

### Accurate posterior inference and conditional sampling in masked VAEs with fixed decoders

As pointed out in the Methods section, the masked training procedure promotes the learning of different encoder networks that share parameters. This allows for targeting different approximate posterior distributions given different conditioning masks: *q*_*ϕ*_(**z**|**x**^all obs^) will most likely not be equal to *q*_*ϕ*_(**z**|**x**^partial obs^), where partial obs indicates that some but not all x-dimensions where observed in contrast to all obs. The generative model (Eq. 1) of the VAE *p*_*θ*_(**x**|**z**), however, should not depend on the masking condition. Once this relationship from latent to data space is correctly capture conditional distributions of unobserved data **x**^unobs^ given observed data **x**^obs^:

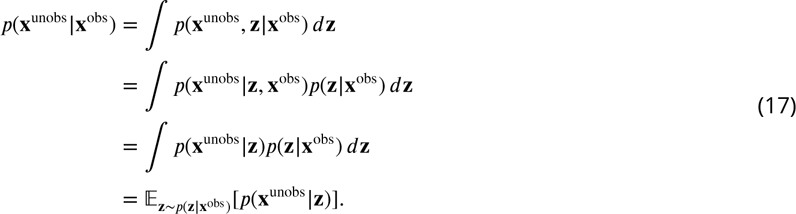

Note that we used that *p_θ_*(**x**∣**z**) can be written as *p_θ_*(**x**^obs^, **x**^unobs^, ∣**z**) and since **x**^obs^ and **x**^unobs^ are conditionally independent given **z**, *p*(**x**^unobs^∣**z**, **x**^obs^) simplifies to *p*(**x**^unobs^ **z**). Hence, assuming we have correctly learned to infer *p*(**z∣x**^obs^) and *p*(**x**^unobs^∣**z**), we can sample from all sorts of conditional distributions we specify prior to training with structured masks.

